## Supplementary figures and images for "Characterization of caffeine response regulatory variants in vascular endothelial cells"

### Figure1_figure_supplement1

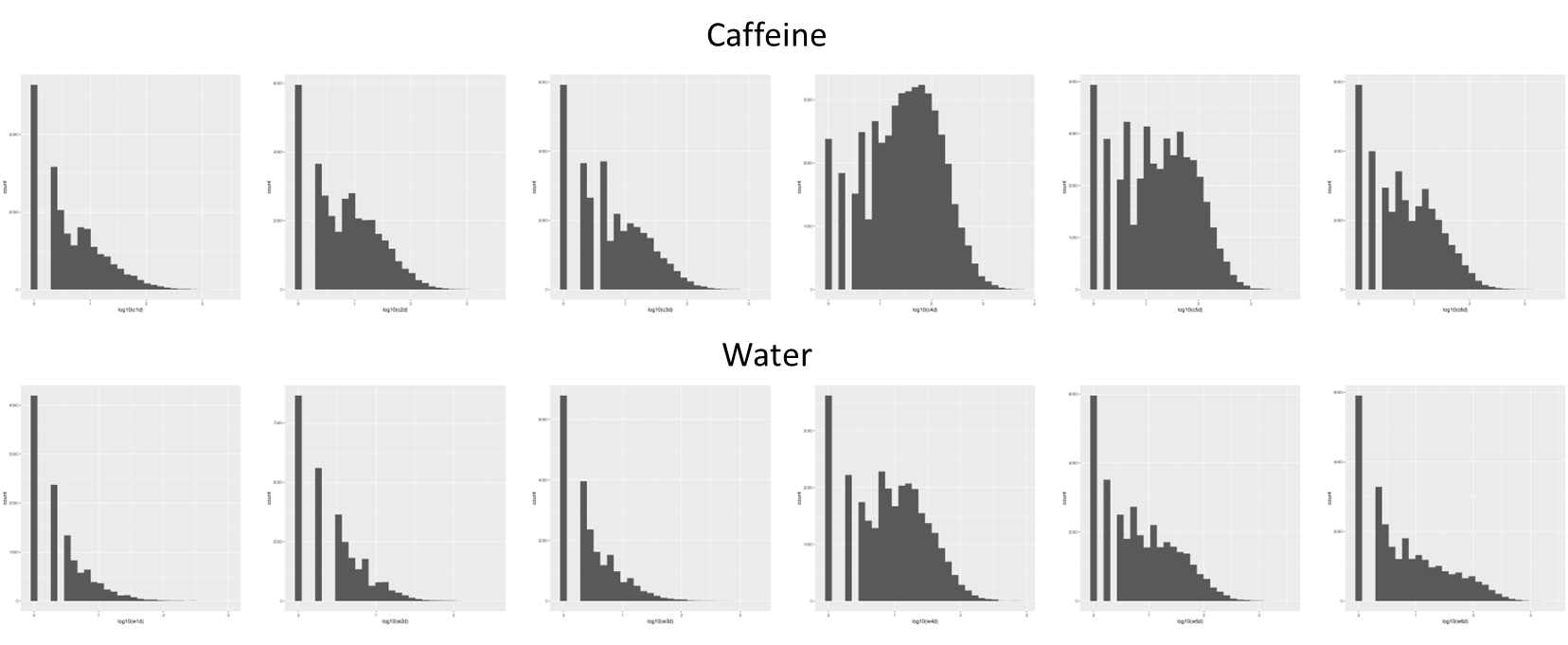

### Figure2_figure_supplement1

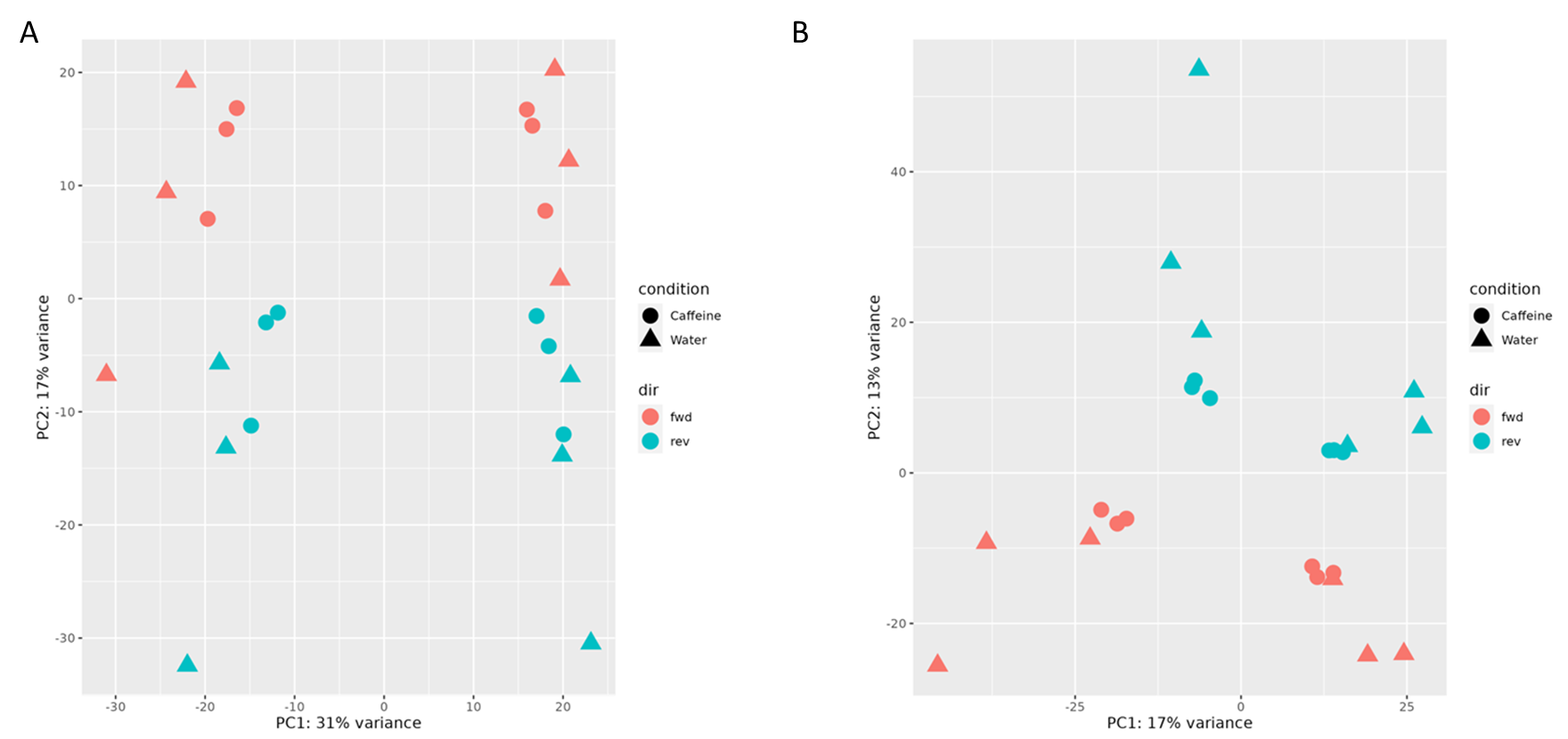

### Figure2_figure_supplement2

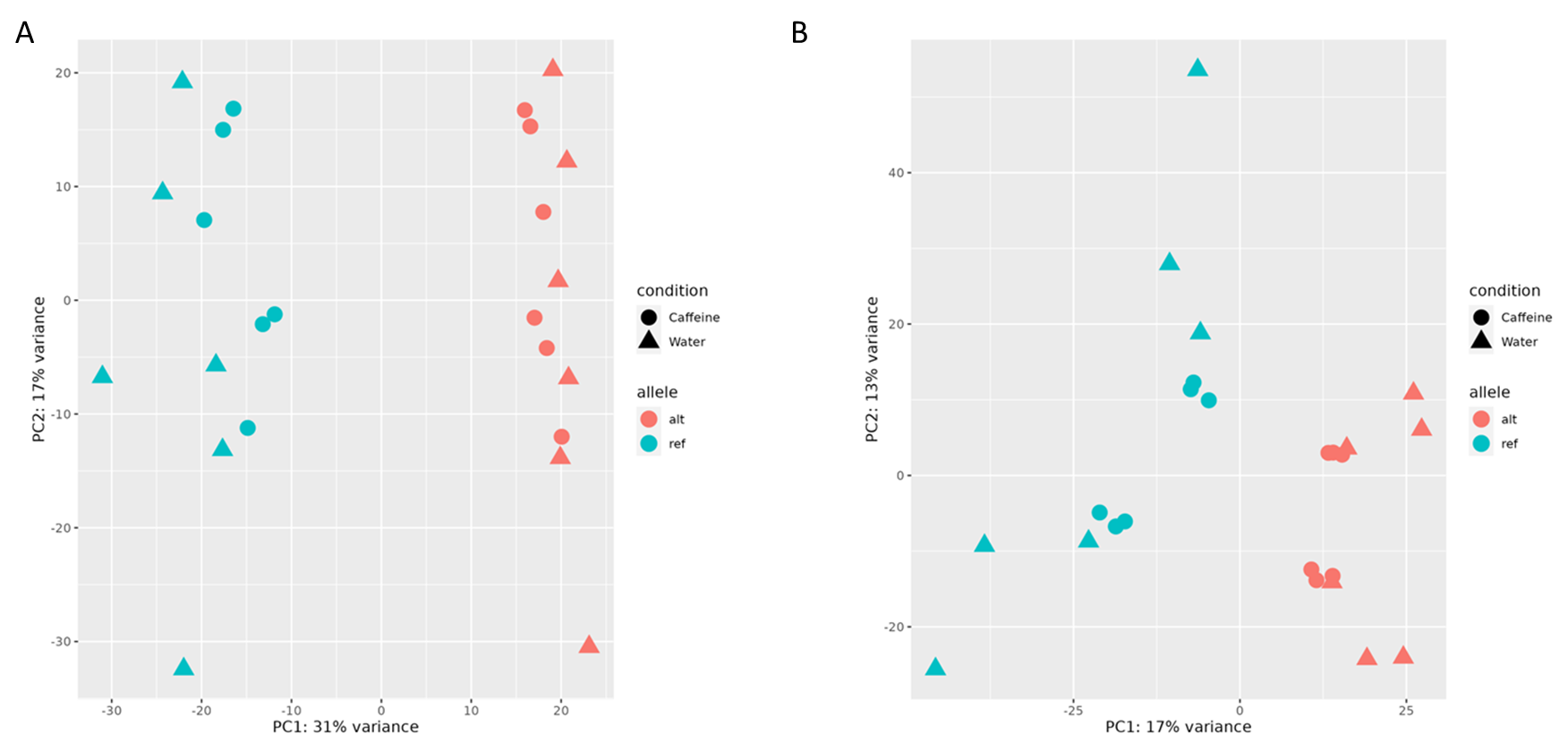

### Figure2_figure_supplement3

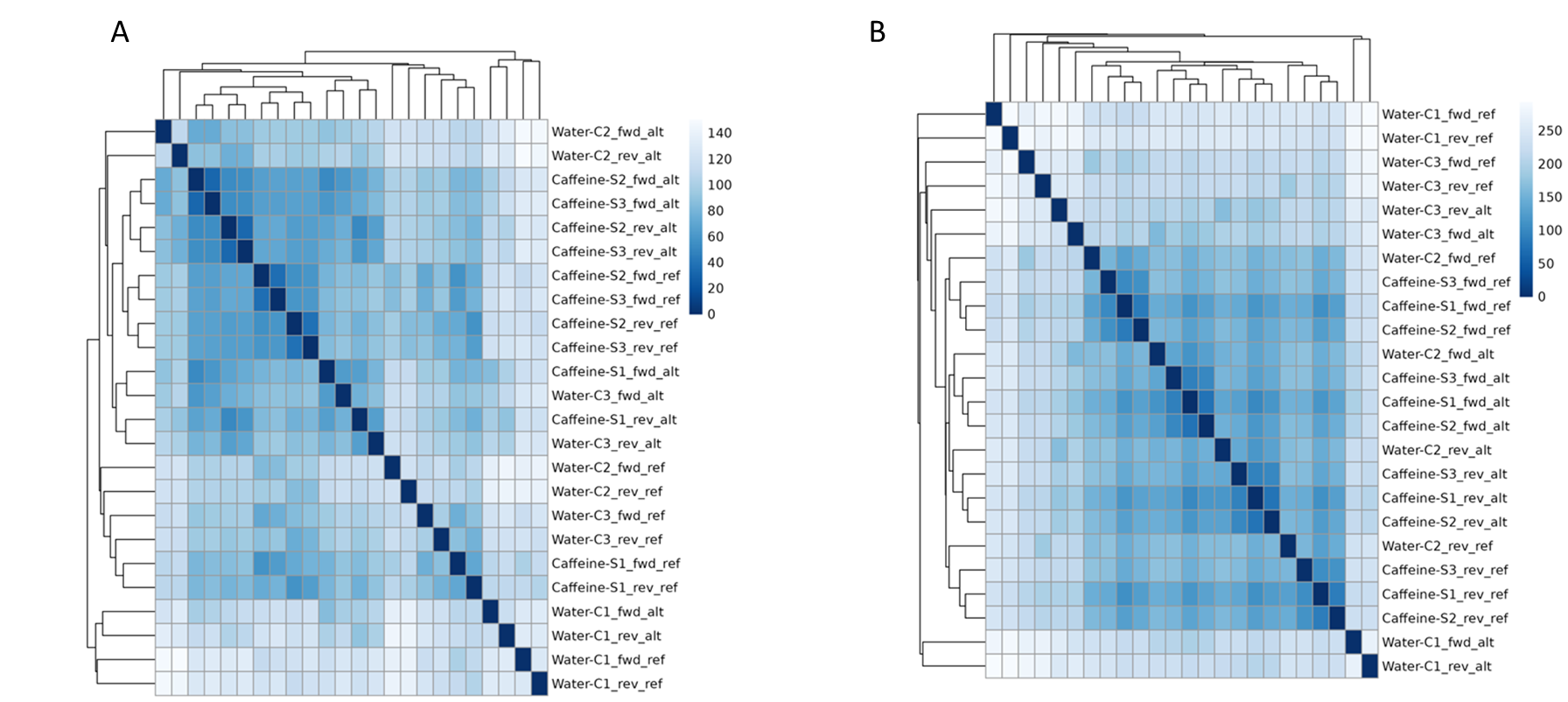

### Figure3_figure_supplement1

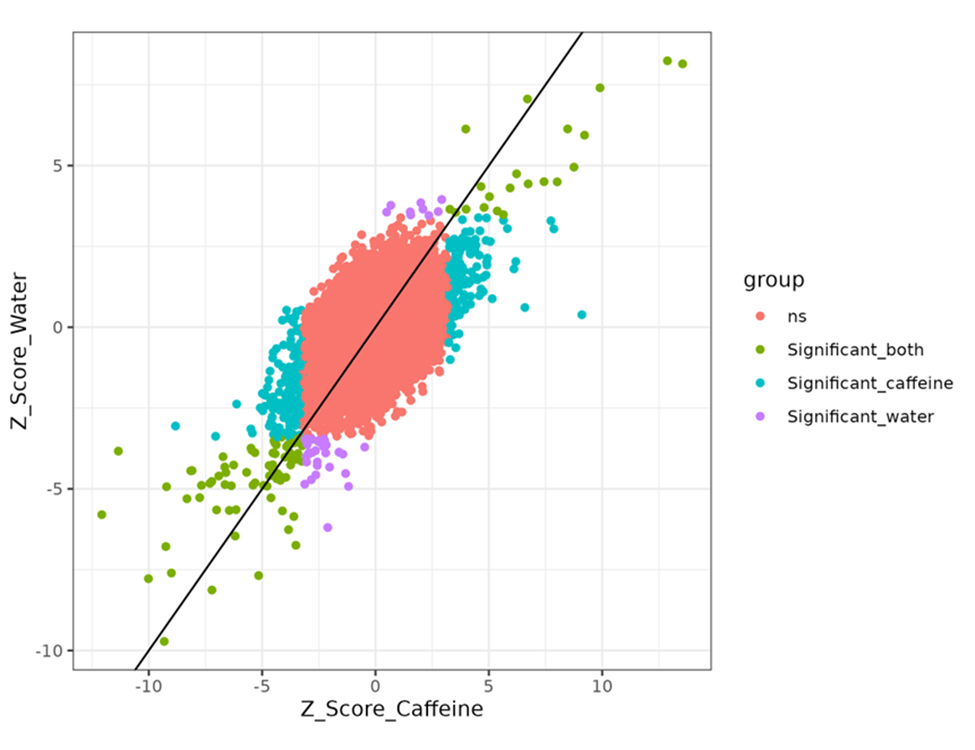

### Figure3_figure_supplement2

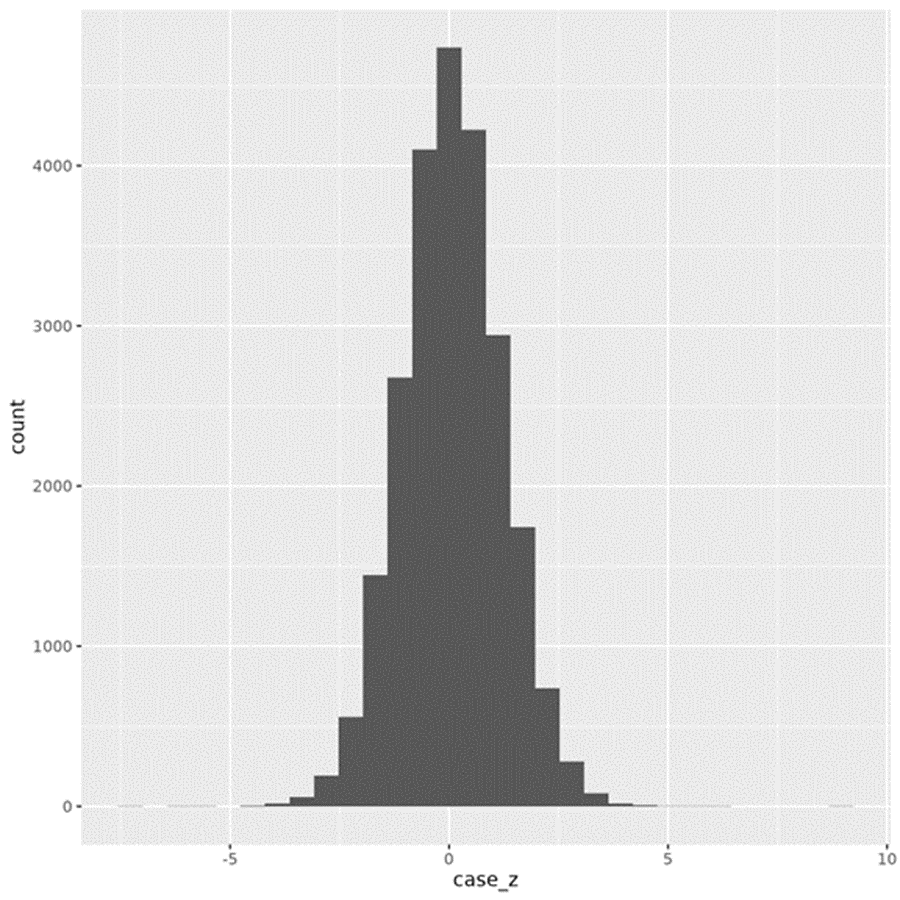

### Figure3_figure_supplement3

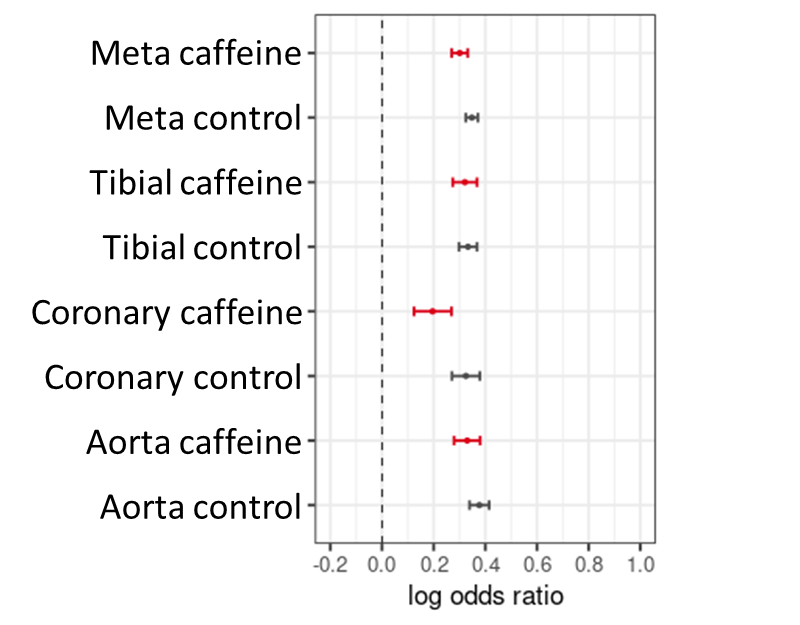
